# Effect of hexanal treatment on fruit qualities and antioxidant activities on ‘Umran’ Indian jujube fruit during cold storage

**DOI:** 10.1101/2023.04.06.535963

**Authors:** Anil Sharma, Harsimrat K. Bons, S K Jawandha, Sun Woo Chung

## Abstract

‘Umran’ Indian jujube is a widely grown cultivar due to its marketable quality and productivity, resulting in excellent market value. Despite its high quality, the product’s short shelf life poses a challenge for transportation, even within the domestic market. Hexanal with three concentrations (0.15%, 0.20%, and 0.25%) was applied to the fruit of ‘Umran’ Indian jujube at both pit hardening and color break stages. Harvested fruits were stored for 28 days in a cold chamber of 7.5±1^°^C and 90-95% RH. The changes in fruit qualities were assessed with physicochemical characteristics, bioactive compounds, and enzymatic antioxidant activities every seven days. Fruits treated with hexanal of 0.20% reduced fruit weight loss and spoilage and maintained fruit firmness, soluble solids content, carotenoid content, and antioxidant activity. The activities of cell wall degrading enzymes investigated were suppressed. Of the antioxidant activities, superoxide dismutase and peroxidase were positively activated. Therefore, these results indicated that pre-harvest application of hexanal 0.20% improved physiochemical characteristics, maintained bioactive compounds, antioxidant enzyme activities, and extended the shelf life of Indian jujube fruits up to 21 days during cold storage.

## Introduction

Indian jujube (*Ziziphus mauritiana* Lamk.) belongs to the *Rhamnaceae* family and is widely grown in arid and subtropical parts of India. Indian jujube has been cultivated mainly in China, Pakistan, Malaysia, and parts of the Gulf countries (Meghwal *et al*., 2022). Jujube fruit is well recognized as a healthy food that contains a variety of bioactive substances, such as polysaccharides, polyphenols, amino acids, nucleotides, fatty acids, dietary fiber, alkaloids, and also contains five times higher vitamin C content than apple fruits and various vitamins including B_1_, B_2_, and P. These nutrients and vitamins possess antioxidant, anti-bacterial, and anti-inflammatory properties, promoting blood circulation, stimulating bile production, and inhibiting various diseases such as allergies and insomnia (Bhargava *et al*., 2005; Zhao *et al*., 2006; Yang *et al*., 2021). Jujube fruits also contain a non-proteinogenic amino acid (aminobutyric acid-GABA) that helps to reduce stress and sleep-enhancing effects.

Numerous cultivars of Indian jujube are commercially grown in India, including ‘Umran’, ‘Gola’, ‘Seb’, ‘Mehroon’, ‘Kaithali’, and ‘Kantha’ (Abdel-Sattar *et al*., 2021). Of these, ‘Umran’ is widely cultivated due to its large size, attractive color, delicious taste, high yield, and rich nutritional value with excellent market value (Azam-Ali *et al*., 2006). Despite their nutritional benefits, ‘Umran’ fruits are susceptible to spoilage, loss of firmness, and reduced shelf-life during transportation and distribution to various regions. The fruits can only last for a maximum of three to five days after harvest before they begin to deteriorate (Chahal and Bal, 2003). These challenges pose a significant threat to the profitability of the fruit industry (Pareek and Yahia, 2013), highlighting the need for innovative and sustainable solutions to improve the shelf-life and transportation of ‘Umran’ fruits.

The postharvest life of fruits is influenced by various factors, including fruit types, storage conditions, and packaging materials (Degl’Innocenti *et al*., 2007; Sajid *et al*., 2019). During postharvest ripening, Indian jujube fruits undergo ethylene evolution, respiratory processes, and changes in cell structures by pectin methyl esterase and polygalacturonase (PG), resulting in fruit softening. These biochemical reactions become challenging to transport ripe fruits to different regions (Goulao *et al*., 2008). To address this challenge, it is necessary to improve the storage life of the fruits through pre-harvest treatments and appropriate storage conditions.

The activity of enzymes and metabolites that degrade the cell wall is closely associated with fruit disorders and the deterioration of fruit quality (Kayal *et al*., 2017). These enzymes include phospholipase D, expansins, pectin methyl esterases (PME), PG, β-galactosidases, and pectate lyases. Fortunately, an appropriate application of hexanal can effectively inhibit the activity of these enzymes, as well as ethylene biosynthesis, metabolic rate, and other enzyme activities that cause fruit spoilage, browning, and degradation of membrane integrity after harvest (Gao *et al*., 2016). Recent studies have demonstrated the effectiveness of hexanal treatment as a pre-harvest spray in extending the shelf life of various fruits, including mango, strawberry, apple, cherry, banana, and blueberry (Paliyath *et al*., 2008). Building on this earlier research, we investigated the effects of different hexanal concentrations on the spoilage, firmness, and postharvest life of Indian jujube fruit.

Hexanal is known for improving the quality and storability of fruits without side effect the fruits (Jincy *et al*., 2017; Ashitha *et al*., 2019). and reported that hexanal treatment reduced fruit browning and spoilage and extended the shelf life of various fruits during cold storage (Erika et al., 2020; Kaur *et al*., 2020). Hexanal treatment in mango fruits improved the shelf life and fruit quality, which would be sociated with improving cell membrane integrity (Preethi *et al*., 2021). Hexanal suppressed the activities of the cell wall-degrading enzymes, including PG, PME, and enzyme phospholipase-D, which is responsible for the deterioration of the fruits storage life (Kumar, 2018). Meanwhile, hexanal promoted the activities of antioxidant enzymes, including catalase (CAT) and peroxidase (POD). This dual effect resulted in decreased rates of specific physiochemical changes in fruits, ultimately improving their storage behaviours. In our study, hexanal was applied to ‘Umran’ Indian jujube fruit, a non-climacteric tropical and arid fruit. The present study aimed to examine the impact of hexanal treatment before harvest to maintain fruit firmness and antioxidant enzyme activities and prevent physiological loss of fruit weight and quality deterioration during cold storage circumstances. Hexanal can inhibit the quality deterioration and antioxidant enzymatic like POD, CAT, super dismutase thereby increasing shelf life and improving the commercial value.

## Materials and Methods

### Plant Materials and Treatments

Fifteen-year-old plants of ‘Umran’ Indian jujube were grown on Fruit Research farm, Punjab Agricultural University, Ludhiana, Punjab (75.791°E, 30.903°N). For treatments, hexanal solutions were prepared with 1% hexanal (v/v), 1% Tween 20 9 (v/v), 1% ascorbic acid (w/v), 1% tocopherol (w/v), 1% geraniol (v/v), and 10 % ethanol 10% (v/v) in distilled water, according to the procedure of Paliyath and Murr, (2007). Three concentrations of hexanal solutions (0.15, 0.20, and 0.25% (v/v)) were applied in mid-December 2021 and early February 2022, which was the color break and pit hardening stages, respectively, with three biological replications.

Fruits were harvested in April at the fully mature stage, precooled, washed, and dried at room temperature. The prepared fruits were wrapped with white paper and placed in corrugated fiber boxes with 5% perforation for cold storage (7.5±1^°^C and 90-95% RH). Thirty fruits were investigated for each biological replication. An experiment was carried out in a completely randomized design with 4 replications, and periodical observation was recorded for physiochemical and enzymatic activities at 0, 7, 14, 21 and 28 days after the cold storage (DACS).

### Determination of physical characteristics

Fruit firmness was determined by using the digital hand penetrometer (FT-327, UC Fruit Firmness Tester, Milan, Italy) with the stainless-steel probe of 8 mm diameter by removing one centimeter peel from two opposite sides of the fruit. The firmness of fruits was expressed in Kg/cm^2^.

Fruit weight loss, physiological loss in weight (PLW), was determined using an equation below:

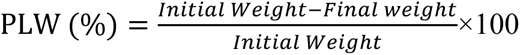

Spoilage was determined by using an equation below:

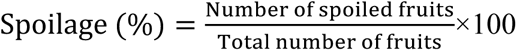

The soluble solids content (SSC) of fruits was determined by using a digital refractometer (RA-250, KEM Kyoto Electronics Manufacturing, Japan) at ambient temperature and expressed in percentage.

Fruit color was determined by using a portable colorimeter (CR-400, Konica Minolta, Japan) which expressed as Hunter Lab (Spectral Range: 400 nm - 700 nm). The instrument measures color in terms of tristimulus value viz. L*, a*, b*. The L*’ value is lightness; the a* value is redness when positive, greenness when negative and b* value is yellowness when positive and blueness when negative.

### Determination of bioactive compounds

#### Ascorbic acid content

By titrating with the dye 2,6-dichlorophenolindophenol (DCPIP), ascorbic acid concentration was determined (AOAC, 2005). For this, 2 ml of juice was mixed with 10 ml of 3% metaphosphoric acid. Then 5 ml of the whole solution was titrated with dye until a rose-pink color appeared.

#### Carotenoid content

Total carotenoids in fruit pulp were determined, according to the procedure of methods proposed by Kirk and Allen, (1965). A 100 mg sample of fruit pulp was finely ground and combined with 2 ml of 80% acetone. Following centrifugation, the resulting supernatant was carefully transferred to a separate tube. The residual pellet was then subjected to an additional extraction using 2 ml of acetone and centrifuged for 10 minutes. Subsequently, both supernatants were combined and diluted with acetone to reach a final volume of 10 ml. Absorbance measurements were taken at three distinct wavelengths: 665 nm, 645 nm, and 480 nm. Carotenoids were calculated by formula below and were expressed as milligrams per 100 grams (mg/100g).

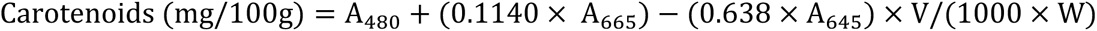

#### Total phenolic content

Folin-Ciocalteu (FC) reagent was used for the estimation as suggested by Hossain and Rahman, (2011). The total phenols were assessed by the technique where juice (5 ml) was taken to 25 ml volumetric flask and add 15 ml of by 80% ethanol for extraction. Then mixture kept for 5 minutes after included of distilled water (6.5 ml) and 1N FC reagent (0.5 ml) in it. To this solution, volume was made to 25 ml by including 1 ml of dehydrated Na_2_CO_3_ and distilled water. This solution were kept for 20 minutes until blue color arose which was perused at 760 nm with spectrophotometer (Theremo Scientific SPECTRONIC 20 D^+^, USA) utilizing gallic acid as a standard and expressed as mg/100 gm pulp.

### Determination of non-enzymatic and enzymatic antioxidant activities

#### Non-enzymatic antioxidant activity

The DPPH (2,2-diphenyl-1-picryl-hidrazil) assay was determined using the method suggested by AOAC, (2005). For estimation of antioxidant activity fruit pulp (3gm) was homogenized in methanol (30 ml) and centrifuged (10,000 × g for 15 minutes at 4 °C). After that, 3 ml (0.1 mM DPPH methanol solution) was added to 100 µL of supernatant, and incubated for 15-20 minutes at 25°C in the dark. The spectrophotometer reading (517 nm) was recorded and total antioxidant activity was calculated by formula:

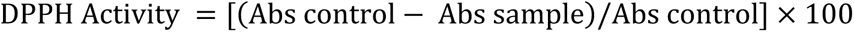

#### Polyphenol oxidase (PPO)

According to Zauberman *et al*., (1991), the 100 mg of fruit pulp was mixed with pH 6.7 phosphate buffer containing 1% of PVP and then centrifuged at 11,000 × *g* for 35 min at 4 °C. By adding 1.0 ml of phosphate buffer (0.1 M) with pH 6.8, 0.1 M 4-methyl catechol (0.5 ml), and 0.5 ml of extraction in a cuvette, the polyphenol activity was measured. The absorbance at 410nm was recorded for 3min in a spectrophotometer (Theremo Scientific SPECTRONIC 20 D^+^, USA). One part of the activity is defined as the amount that results in a 0.01 rise in absorbance per minute and expressed as units.min ^-1^g FW.

#### Peroxidase (POD)

According to the procedure of Shannon *et al*., (1665), the 100 mg of fruit pulp was mixed with pH 6.5 buffer containing 1% of PVP and centrifuged (10,000 ×g for 30 min; 4 °C). For the enzymatic activity, the supernatant was used for assays after the contents were transferred to a spectrophotometric cuvette and the reaction was initiated by 0.1 ml of hydrogen peroxide directly in the cuvette. In comparison to the blank reagent, absorbance was measured at 430 nm every 30 seconds, and activity was expressed as units.min ^-1^g FW.

#### Pectin methyl esterase (PME)

According to Mahadevan and Sridhar, (1882), the 20 mg of pulp was taken and mixed with NaCl solution (80 ml) and centrifuged at 448× g for 30 min at 4℃ after that filtered. The supernatant extract was used to assay the PME enzymatic activity and was expressed as µmol of methyl ester gm^-1^ FW min ^-1^.

#### Polygalacturonase (PG)

By using the method of Lohani *et al*., (2004), with minute moderation and ber pulp was blended for 30 min in 0.5 Tris-HCl buffer containing 1 mM EDTA (5% PVP). After that centrifugation at 10,000 × *g* for 30 min at 4°C. The enzyme activity was determined by the lucid supernatant and expressed as ug of D-galactose gm^-1^ FW min ^-1^.

#### Catalase (CAT)

According to Hu *et al*., (2014), the CAT activity was determined by using a 100 mg sample taken from 4 to 5 fruits pulp homogenized with phosphate buffer (pH 7.5) on chilled ice, and centrifuged at 11,000 × *g* for 30 min at 4°C. The enzyme activity was determined and activity expressed as units.min ^-1^g FW.

#### Superoxide dismutase (SOD)

By the method suggested by Zhang *et al*., (2010), the SOD activity was determined by 100 mg fruit pulp taken from 3 to 5 fruit and mixed with potassium phosphate buffer having pH 7.2 containing 1mM EDTA (1% PVP), 10 mM b-mercaptoethanol on chilled ice, and centrifuged for 11,000 × *g* for 25-30 min at 4°C. The supernatant was used for the assay of the enzyme activity by spectrophotometric at 420 nm up to 3 minutes and was expressed as units.min ^-1^g FW.

### Statistical Analysis

Statistical analysis was performed with Pearson’s correlation analysis, the analysis of variance, and least significant difference test with *p* ≤ 0.05. on S.A.S. Software (Version 9.3). The experimental data was displayed as mean with standard deviation.

## Results

### Changes in fruit physicochemicals

#### Fruit Firmness and spoilage

The hexanal treatment demonstrated an inhibitory effect on abrupt changes in fruit firmness (Table 1), effectively preserving the quality and freshness of Indian jujube fruit. Fruit firmness initially decreased and subsequently increased during cold storage. After 28 days after cold storage (DACS), 0.20% hexanal-treated fruit maintained a fruit firmness of 0.07 Kg/cm2, indicating a 20.35% higher fruit firmness compared to untreated fruits. Hexanal-treated fruits exhibited improved firmness compared to control fruits. The current findings (Table 2 and Fig. 3a) revealed a negative correlation (−0.962, R² = 0.8312) between firmness and PLW. Higher PLW during fruit storage is associated with decreased firmness and reduced turgidity due to the loss of moisture content in the fruit. Among all the treatments, no fruit spoilage was observed up to 7 days after cold storage (DACS) (Table 1). After 14 DACS, spoilage of untreated fruit reached 2.56%, while hexanal-treated fruits remained unchanged. The spoilage of hexanal-treated fruits was lower up to 21 DACS compared to untreated fruits. After 28 DACS, untreated fruits exhibited higher spoilage (37.45%) than treated fruits, and 0.20% hexanal-treated fruits had the lowest spoilage (23.54%). Fruit spoilage displayed negative correlation coefficients with fruit firmness, corresponding to changes in firmness (Table 2).

**Table 1.** Variation in different parameters like physiological loss in fruit weight, spoilage and fruit firmness of Indian jujube fruits cv. Umran under cold storage about different concentration of hexanal under cold storage conditions.

| Variable | Hexanal treatment | Storage period (days after cold storage) |  |  |  |  |
| --- | --- | --- | --- | --- | --- | --- |
|  |  | 0 | 7 | 14 | 21 | 28 |
| Fruit Firmness (Kg/cm<sup>2</sup>) | 0.15% | 14.75 ±0.03<sup>b</sup> | 14.24 ±0.03<sup>b</sup> | 12.34 ±0.04<sup>b</sup> | 11.24 ±0.04<sup>b</sup> | 9.92 ±0.01<sup>b</sup> |
|  | 0.20% | 14.85 ±0.02<sup>a</sup> | 15.55 ±0.04<sup>a</sup> | 12.40 ±0.03<sup>a</sup> | 11.54 ±0.06<sup>a</sup> | 10.07 ±0.02<sup>a</sup> |
|  | 0.25% | 14.55 ±0.06<sup>c</sup> | 14.10 ±0.05<sup>c</sup> | 12.00 ±0.02<sup>c</sup> | 11.23 ±0.05<sup>b</sup> | 9.95 ±0.01<sup>b</sup> |
|  | Control | 13.36 ±0.04<sup>d</sup> | 13.11 ±0.04<sup>d</sup> | 11.9 ±0.07<sup>d</sup> | 10.06 ±0.06<sup>c</sup> | 8.02 ±0.03<sup>c</sup> |
| PLW (%) | 0.15% | — | 1.20 ±0.035<sup>c</sup> | 1.57 ± 0.04<sup>bc</sup> | 2.51 ±0.05<sup>b</sup> | 4.48 ±0.05<sup>b</sup> |
|  | 0.20% | — | 1.15 ±0.03<sup>d</sup> | 1.36 ±0.05<sup>c</sup> | 2.44 ±0.02<sup>c</sup> | 4.38 ±0.04<sup>c</sup> |
|  | 0.25% | — | 1.30 ±0.06<sup>b</sup> | 1.53 ±0.07<sup>b</sup> | 2.47 ±0.06<sup>b</sup> | 4.41 ±0.06<sup>b</sup> |
|  | Control | — | 2.60 ± 0.05<sup>a</sup> | 3.93 ±0.05<sup>a</sup> | 4.25 ±0.03<sup>a</sup> | 6.63 ±0.07<sup>a</sup> |
| Spoilage (%) | 0.15% | — | — | 0.00 ±0.00<sup>b</sup> | 4.34 ±0.07<sup>b</sup> | 26.36 ±0.17<sup>b</sup> |
|  | 0.20% | — | — | 0.00 ±0.00<sup>b</sup> | 3.90 ±0.06<sup>c</sup> | 23.54 ±0.15<sup>d</sup> |
|  | 0.25% | — | — | 0.00 ±0.00<sup>b</sup> | 4.23 ±0.08<sup>b</sup> | 24.54 ±0.11<sup>c</sup> |
|  | Control | — | — | 2.56 ±0.08<sup>a</sup> | 9.14 ±0.08<sup>a</sup> | 37.45 ±0.18<sup>a</sup> |

Mean values followed by the different superscript within a column are significantly different at $p \leq 0.05$ .
PLW, physiological loss in weight.

**Table 2.** Pearson’s correlation coefficients between variables of ‘Umran’ Indian jujube fruits.

| Variable | PLW | Spoilage | SSC | Firmness | Acidity | AA | SOD | POD | Catalase | PME | PPO |
| --- | --- | --- | --- | --- | --- | --- | --- | --- | --- | --- | --- |
| Spoilage | 0.996** | 1 |  |  |  |  |  |  |  |  |  |
| SSC | -0.035* | 0.041* | 1 |  |  |  |  |  |  |  |  |
| Firmness | -0.962* | -0.976* | -0.070** | 1 |  |  |  |  |  |  |  |
| Acidity | 0.385* | 0.319* | -0.925** | -0.309** | 1 |  |  |  |  |  |  |
| AA | 0.229* | 0.144* | -0.856** | -0.012* | 0.797* | 1 |  |  |  |  |  |
| SOD | -0.872* | -0.911* | -0.428** | 0.928** | 0.061** | 0.266* | 1 |  |  |  |  |
| POD | -0.525* | -0.550* | -0.610** | 0.428** | 0.450** | 0.140* | 0.656** | 1 |  |  |  |
| Catalase | 0.083* | -0.001* | -0.959** | 0.089** | 0.867** | 0.963* | 0.412** | 0.400** | 1 |  |  |
| PME | -0.913* | -0.876* | -0.429** | 0.815** | -0.702** | -0.588* | 0.600** | 0.281** | -0.480** | 1 |  |
| PPO | 0.149* | 0.211* | 0.328** | -0.412** | -0.119** | -0.692* | -0.434** | 0.243** | -0.564** | 0.077** | 1 |
| PG | 0.970* | 0.978* | -0.008** | -0.996** | 0.383** | 0.0867* | -0.898** | -0.390** | -0.010** | -0.855** | 0.376** |

\*, \*\*Significant at $p \leq 0.05$ and $p \leq 0.01$ , respectively.
AA, antioxidant activity.

#### Fruit weight loss

Higher weight loss observed in stored fruits was associated with their firmness and turgidity (Table 2). Throughout cold storage, the PLW of the fruits consistently increased (Table 1). Hexanal treatments significantly reduced weight loss in fruits when compared to untreated fruits during cold storage. At 28 DACS, the maximum fruit weight loss (6.63%) was noted in untreated fruits, whereas fruits treated with 0.20% hexanal demonstrated the lowest weight loss (4.38%).

#### Fruit color values (L*, a*, b*)

During cold storage, the fruit peel colors changed (Fig. 1d-f). As shown in Fig. 1d, the fruit color L* value decreased up to 21 DACS. At 28 DACS, the minimum and maximum L* values (31.54 and 37.03) were observed in 0.20% hexanal-treated fruits and untreated fruits, respectively. The a* value was highest (2.39) in untreated fruits and lowest (1.73) in 0.25% hexanal-treated fruits (Fig. 1d). The fruit color b* value was lowest (29.99) in 0.20% hexanal-treated fruits after 14 DACS, and then declined to 17.86. The maximum value (21.56) was found in untreated fruits after 28 DACS (Fig. 1f).

**Fig. 1.**
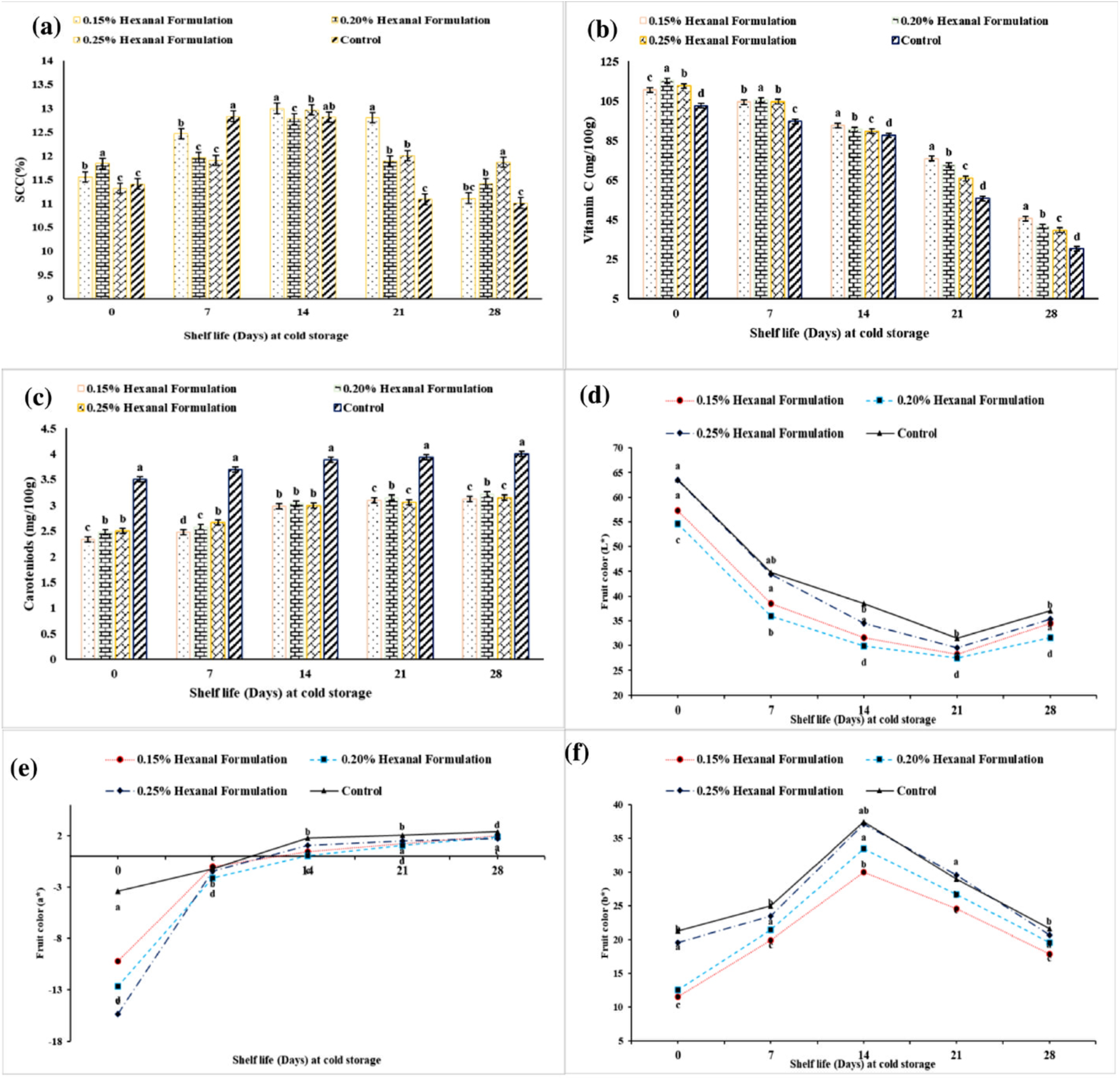
Variation in soluble solids content, ascorbic acid content, carotenoids, and color values (L*, a*, b*) of ‘Umran’ Indian jujube fruits during cold storage by different hexanal treatments. Mean values depicting the different superscript within days after cold storage are significantly different at *p* ≤ 0.05.

### Soluble solids content

The SSC initially increased and then decreased up to 28 DACS (Fig. 1a). The SSC continuously increased until 14 DACS, reaching 13.00% and 12.97% in fruits treated with 0.15% and 0.25% hexanals, respectively. The SSC of untreated fruit was higher at 7 DACS and declined by 28 DACS. The SSC exhibited a negative correlation coefficient with firmness, indicating that a decrease in firmness would be associated with an increase in SSC (Table 2).

### Changes in bioactive compounds and antioxidant activities

For all treatments, the ascorbic acid content declined up to 28 DACS (Fig. 1b). The reduction in ascorbic acid content was less pronounced in hexanal-treated fruits compared to the control treatment. Fruits treated with 0.20% hexanal initially had the highest ascorbic acid content (105.43 mg). However, after 28 DACS, the maximum content (41.56 mg) was observed in fruits treated with 0.15% hexanal, as opposed to the untreated fruits.

Hexanal treatments influenced carotenoid content in Indian jujube fruits under cold storage, as shown in Fig. 1c. Carotenoids increased steadily as the storage intervals progressed. The interaction between storage intervals and different hexanal treatments was found to be statistically significant. After 28 DACS, the minimum carotenoid content (3.12 mg) was found in fruits treated with 0.15% hexanal, while the maximum content (4.00 mg) was observed in untreated fruits.

The phenol content and antioxidant activity decreased throughout the storage period, regardless of the treatments. The levels of antioxidant activity and phenols were significantly higher in hexanal-treated fruits, as shown in Fig. 2(a-b). At 0 DACS, no significant difference was observed. After 7 DACS, the total antioxidant activity and phenols gradually decreased up to 28 DACS for all treatments. At the end of the experiment, the maximum antioxidant activity (13.61 µmol/g) and phenols (100.27 mg) were observed in fruits treated with 0.20% and 0.15% hexanal, respectively, compared to untreated fruits.

**Fig. 2.**
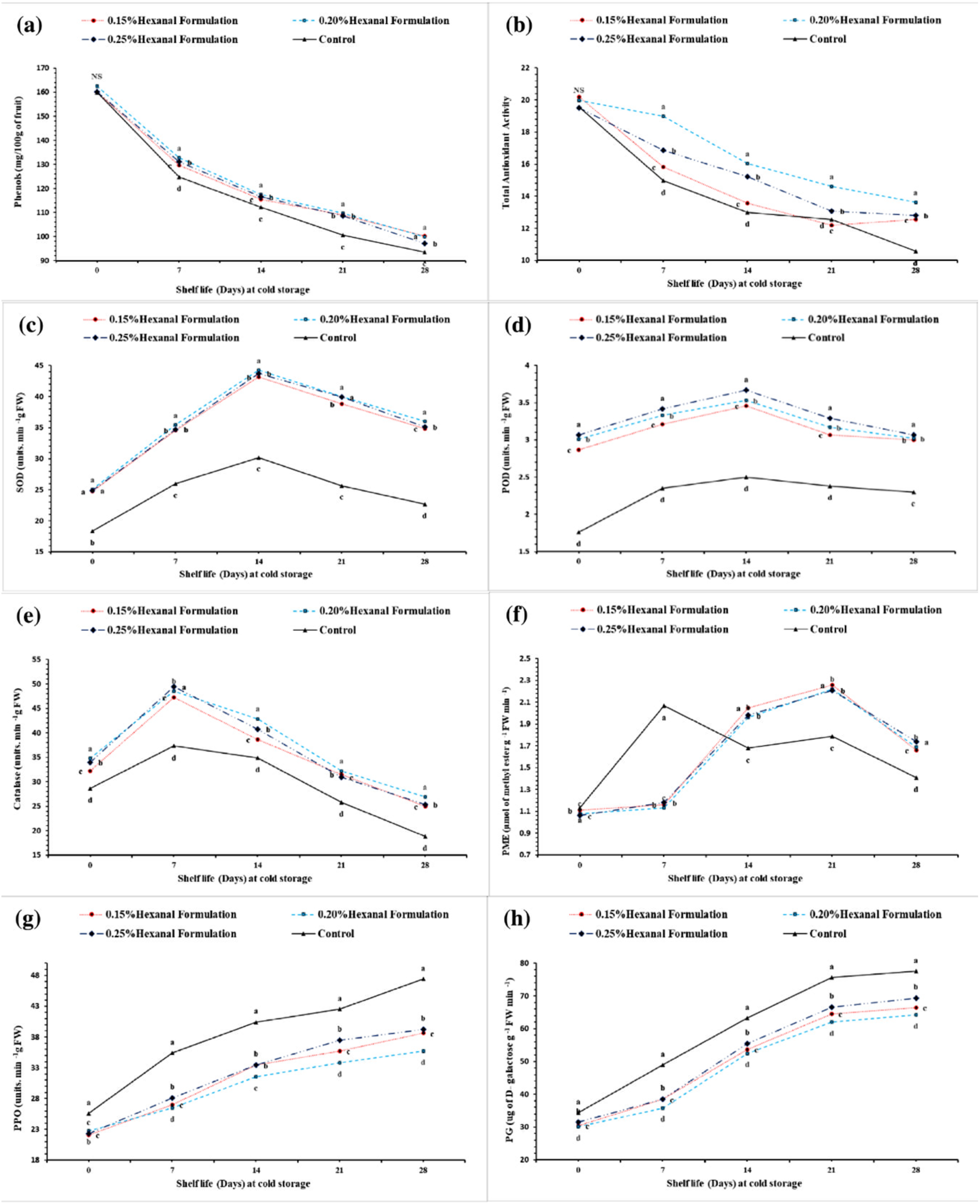
Variation in phenol content, and non-enzymatic and enzymatic antioxidant activities, including superoxide dismutase (SOD), peroxidase (POD), catalase (CAT), pectin methyl esterase (PME), polyphenol oxidase (PPO) and polygalacturonase (PG). Vertical bars represent S.D. of means for four replicates. Mean values depicting the different superscript within days after cold storage are significantly different at *p* ≤ 0.05.

The SOD activity for different treatments was depicted in Fig. 2c. The SOD activity increased as storage progressed up to 14 DACS, followed by a decline at 28 DACS. The SOD activity of the control fruit was consistently lower than that of other treatments throughout the storage period. In contrast, the SOD activity of fruits treated with hexanal exhibited consistently higher activity. In the end, fruits treated with 0.20% hexanal had higher SOD activity (37.11%) compared to control fruits. Correlation coefficients showed significant (p ≤ 0.05) results between SOD and spoilage (−0.428) during the cold storage conditions (Table 2).

The POD activity varied throughout the cold storage period, irrespective of the given treatments, as shown in Fig. 2d. At the beginning of storage, the POD activity slightly increased, reaching a maximum on the 14 DACS, which was higher (29.17%) than the control. After that, the POD activity decreased. After 28 DACS, the enzyme activity of the untreated fruits was the lowest, while that of 0.20% hexanal-treated was fruits was the highest. Pearson correlation coefficient showed a significant correlation between PME activity and physiological loss in weight (−0.525) during the storage period (Table 2).

The CAT activity initially increased with the progression of the storage period up to 7 DACS and declined afterward (Fig. 2e). Irrespective of all treatments, after 7 DACS, 0.20% hexanal-treated fruits reached higher (24.11%) activity than untreated fruits, and then it declined abruptly. At the end of storage, the maximum (36.05 units.min-1g FW) enzyme activity was recorded in 0.20% hexanal-treated fruits. The firmness of Indian jujube fruits was negatively correlated with CAT (−0.959, R^2^ = 0.5633) during the storage period (Table 2 and Fig. 3d).

**Fig. 3.**
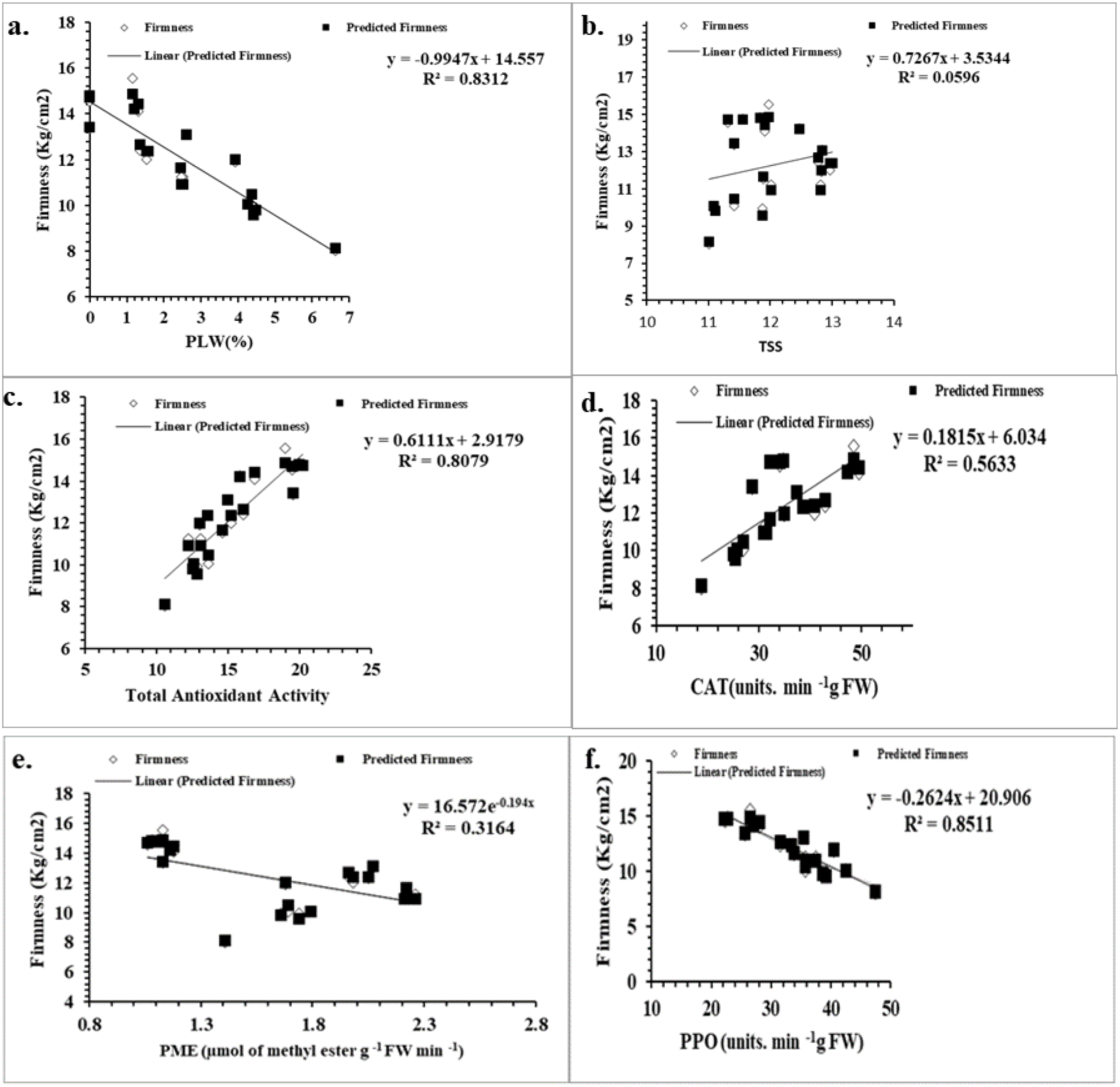
Regression relation between firmness and polyphenol oxidase (PPO), catalase (CAT), pectin methyl esterase (PME), total antioxidant activity (AoA), physiological loss in fruit weight (PLW), and soluble solids content (SSC).

The PME activity significantly varied during the storage of fruits. Initially, it increased up to 21 DACS, and later a decline in activity was observed on the 28 DACS (Fig. 2f). In the beginning, untreated fruits showed maximum enzyme activity compared to treated fruits. The fruits treated with 0.15% hexanal reached 20.80% higher enzyme activity than untreated fruits on the 21 DACS, and afterward, it declined in all the treatments. Fruit firmness was inversely correlated with PME cell wall-degrading enzyme activity during storage (Table 2 and Fig. 3e). PME and firmness were found to have a significant (p ≤ 0.05) Pearson correlation coefficient (−0.429, R2 =0.3164) during the storage period; an increase in PME activity led to a reduction in fruit firmness (Table 2 and Fig. 3e).

The PPO activity of all the treatments increased progressively during cold storage, as illustrated in Fig. 2g. However, the polyphenol activity of untreated fruits was significantly higher (24.73%) compared to treated fruits after 28 DACS. At the end of the experiment, 0.20% hexanal-treated fruits recorded the lowest enzyme activity. The results revealed that the Pearson correlation coefficient showed a significant (p ≤ 0.05) correlation between PPO with SSC (−0.412) and PPO with Firmness (0.328, R2 =0.8511) during the storage (Table 2 and Fig. 3f).

The PG activity initially increased with the progression of the storage period throughout the experiment in all the treatments (Fig. 2h). After 28 DACS, untreated fruits showed 17.20% higher PG activity than fruits treated with 0.20% hexanal and also recorded significant results in Pearson correlation coefficient relation between PG and firmness (−0.008) during the cold storage condition (Table 2).

## Discussion

Perishable fruits like Indian jujube have a limited shelf life due to various internal and external factors [Pareek and Yahia, 2013). PLW in fruits typically occurs because of transpiration, respiration, and other metabolic processes during storage, leading to cell wall breakdown and gas exchange between the inner and outer tissues of the fruit membrane (Ramezanian *et al*., 2010). Under standard storage conditions, starch rapidly converts to sugars in fruits, and the degradation of organic acids and polyphenols occurs, making them susceptible to microorganisms such as fungi and bacteria, which can cause postharvest diseases (Caruso and Ramsdell, 1995).

Hexanal is a chemical known to effectively inhibit the phospholipase-D enzyme, thereby extending the storage life of fruit crops. The activity of phospholipase-D during ripening, along with reactive oxygen species generated during enzymatic activity and under stress, promotes senescence in fruit crops (Paliyath *et al*., 2008). During cold storage, hexanal reduces physiological weight loss by serving as a barrier for gas exchange and water loss (Hoa and Ducamp, 2005). Weight loss occurs during ripening due to the significant energy required for the process, as starch is converted into sugar and used as an energy source (Ashwini *et al*., 2018).

Hexanal has also proven effective in preventing fungal growth during storage, inhibiting spore production, reducing fruit spoilage and decay incidence, and increasing PME activity, which helps maintain fruit quality by regulating enzymatic activities (Baggio *et al*., 2014; Gill *et al*., 2016). Hexanal has demonstrated potential in improving shelf life and biochemical properties of various fruits, including apples, bananas, cherries, strawberries, and pears (Paliyath *et al*., 2008). Hexanal treatment leads to decreased transcriptional expression of PG and PME, which in turn slows pectin breakdown, increases fruit firmness, and preserves cell membrane integrity during cold storage.

Physical parameters of fruits, such as firmness, spoilage, color, and texture, are crucial factors that determine the quality, aroma, and acceptability of fruits, ultimately enhancing their table value and industrial applications (Salehi, 2021). The primary textural changes responsible for fruit softening result from enzyme-mediated alterations in the cell wall structure and pectin content, specifically the solubilization of the polysaccharides composing the cell wall (pectins and celluloses) either entirely or partially (Jha *et al*., 2006; Reddy *et al*., 2017). Rapid ripening in untreated fruits during storage leads to a more pronounced decline in total phenol and pectin content compared to treated fruits, which aligns with observations made for ber fruits (Singh and Bal, 2006) and cherry fruits (Sharma *et al*., 2006) during storage.

Increased fruit weight loss and decreased fruit firmness may be attributed to the promotion of cell wall breakdown by PG and PME. PME catalyzes the de-esterification of pectin, producing short de-methylated pectin chains that are more susceptible to pectin depolymerase, ultimately causing significant changes in fruit cell walls and softening of fruits (Anusuya *et al*., 2016; Gill *et al*., 2016). Hexanal treatment leads to reduced depolymerase and PME activity, which decreases pectin content, extends shelf life, and preserves fruit firmness (Kumari *et al*., 2017).

During cold storage, PG catalyzes PME activity, accelerating pectin depolymerization through pectin degradation and fruit softening (Gill *et al*., 2016; Kaur *et al*., 2020). Positively charged aliphatic amines (polyamines) strengthen cell walls by forming cross-links between carboxyl groups, which slow down the activity of enzymes like PG and PME during storage and delay cell wall weakening (Valero *et al*., 2002). Similar to our findings, previous research has shown that pear softening is also influenced by polyamines (Singh *et al*., 2019), and a slowdown in enzymatic activities, such as PG and PME, occurs in peach fruits (Torrigiani *et al*., 2012). Hexanal spray effectively reduced PME activity, respiration rate, and fruit decay incidence. Hexanal-treated mango fruits exhibited increased soluble solids content, firmness, and fruit acceptability up to 28-30 days of storage (Kaur *et al*., 2020). In this study, a clear negative correlation was observed between PG and PME activities in relation to fruit firmness, ultimately impacting fruit freshness and storage life.

Fruit color changes are favorably associated with total carotenoid content in Indian jujube fruits (Shi *et al*., 2018). Fruit color positively correlates with total carotenoids, which increase with fruit ripening (Mercadante *et al*., 1998). Indian jujube fruits exposed to hexanal spray exhibit color development and retain their green color longer than untreated fruits (Anusuya *et al*., 2016). The present study demonstrated a positive correlation between carotenoid synthesis and hexanal treatment, helping maintain fruit color for up to 21 days after cold storage (DACS).

Hexanal treatment inhibits POD activity by indirectly preventing free radical production (Kaur *et al*., 2020). Compared to fruits stored at low temperatures, the change in POD activity is more pronounced in fruits held at ambient temperatures, possibly due to the temperature-dependent nature of the POD enzyme, which is adversely affected by increasing temperatures (Yingsanga *et al*., 2008). Hexanal treatment enhances fruit freshness, chroma value, color, firmness, anthocyanins, and phenolic components while increasing SOD and POD activities (Habibi *et al*., 2022). In the current study, pre-harvest hexanal treatment regulated enzymatic activities and maintained firmness compared to control fruits throughout the storage period, effectively extending the shelf life of Indian jujube fruits under cold storage conditions.

## Conclusion

Our study demonstrated that hexanal treatment effectively preserves postharvest quality and antioxidant enzymatic activities in Indian jujube fruits. The application of 0.20% hexanal reduced spoilage rates, inhibited weight loss, and maintained fruit firmness for longer durations. Our results indicate that pre-harvest hexanal application has considerable potential to suppress ethylene production, acting as an efficient method for delaying ripening and maintaining the quality and storability of Indian jujube fruits during cold storage. The 0.20% hexanal treatment can effectively maintained fruit firmness by reducing cell wall-degrading enzymatic activities, such as polyphenol oxidase, PG, and POD, under cold storage conditions. Additionally, these treatments aid in preserving fruit qualities, including soluble solids content, ascorbic acid, fruit color (L*, a*, b*), and carotenoid contents. Consequently, pre-harvest application serves as an effective measure to maintain high quality and extend the postharvest shelf life of Indian jujube fruits for up to 21 days under cold storage conditions.

## Acknowledgements

The authors are thankful to the Punjab Agricultural University, Ludhiana (Punjab), India, for providing the essential research facilities.

## Declaration

### Data availability

All data generated or analysed during this study are included in this article.

### Authors’ contributions

AS: Writing-original draft, Formal analysis, Investigation, Methodology, Data curation, Software, Validation HKB: Supervision, Conceptualization, Investigation, Methodology, Visualization, Resources SKJ: Supervision, Data curation, Formal analysis, Investigation, Visualization SWC: Writing-review & editing. Methodology, Conceptualization

